# Translational approach to increase phosphate accumulation in two plant species through perturbance of inositol pyrophosphates

**DOI:** 10.1101/2022.05.02.489396

**Authors:** Catherine Freed, Branch Craige, Caitlin Cridland, Janet Donahue, Sarah Phoebe Williams, Jiwoo Kim, Glenda Gillaspy

## Abstract

Inorganic phosphate (P*i*), while indispensable for all biological organisms and a major agricultural macronutrient, is an increasingly limited and nonrenewable resource. Recent studies demonstrate the importance of inositol pyrophosphates (PP-InsPs) in plant P*i* signaling and homeostasis, however the extent to which PP-InsPs impact plant development is not well understood. We report that transgenic expression of the *Saccharomyces cerevisiae* enzyme Diadenosine and Diphosphoinositol Polyphosphate Phosphohydrolase (DDP1) in *Arabidopsis thaliana* and *Thlaspi arvense* (pennycress) provide a unique translational utility for P*i* phytoremediation as well as unique germplasm and insight on the long-term impacts of reduced PP-InsPs. Transgenic DDP1 expression in *Arabidopsis* decreased PP-InsPs, impacted growth and development, and increased P*i* accumulation leading to P*i* toxicity. Analysis of P*i* Starvation Response (PSR) marker genes indicated that the PSR is activated in DDP1 expressing plants. We assessed translational utility through transformation of pennycress, a spring annual cover crop with emerging importance as a biofuel crop, with a *DDP1* transgene. Pennycress plants expressing DDP1 showed similar altered P*i* accumulation phenotypes, suggesting that these plants could potentially serve to remove P*i* from P*i*-rich soils. Our study addresses the long-term impacts of PP-InsP reduction on plant growth, as well as establishing a starting material for a unique P*i* reclaiming cover crop.

**SIGNIFICANCE STATEMENT:** A major challenge to food security is the phosphorus (P) crisis. A global P shortage is imminent based on the misuse of current resources and will be further aggravated by climate change and a lack of policy addressing sustainability. Our work addresses this crisis by investigating the sustained impact of altering inositol pyrophosphates to manipulate plant P accumulation, a strategy that could be used to remediate nutrient-polluted environments.

## INTRODUCTION

Inorganic phosphate (P*i*), a required nutrient for genetic maintenance, cellular function, and energy metabolism, is arguably the greatest plant growth-limiting macronutrient and is indispensable for food production and security on a global scale (Vance *et al*., 2003). While critically important, P*i* is scarce in many agricultural soils and is a rapidly depleting, nonrenewable resource (Cordell *et al*., 2009). Moreover, current P*i* usage is largely inefficient and surplus application of P*i* leads to environmental pollution (Elser and Bennett, 2011). A worldwide P shortage, also referred to as the P crisis (Abelson, 1999; Cordell *et al*., 2009), is imminent and is predicted to detrimentally impact global food security. Given these challenges, it is crucial to understand how plants utilize P*i* to enhance efficiency of P usage. Under depleted P*i* conditions, plants employ molecular mechanisms, collectively known as the P*i* starvation response (PSR), that reprioritize growth patterns to increase P*i* uptake, and redistribute P*i* from existing cells (Rouached *et al*., 2010; Chien *et al*., 2018; Wang *et al*., 2021). The PSR is modulated by complex signaling networks and while its regulation is not completely understood, emerging evidence strongly supports an integral role for inositol pyrophosphates (PP-InsPs) as regulators of the PSR *in vivo* (Wild *et al*., 2016; Dong *et al*., 2019; Zhu *et al*., 2019).

PP-InsPs and their precursors, inositol phosphates (InsPs), are crucial for development, energy metabolism, and stress responses in plants (Tsui and York, 2010; Williams *et al*., 2015; Freed *et al*., 2020). InsPs consist of a *myo*-inositol ring with hydroxyl groups that are sequentially phosphorylated by evolutionarily conserved enzymes (Irvine and Schell, 2001; Williams *et al*., 2015; Livermore *et al*., 2016). The number and position of phosphate moieties on the ring convey different intracellular messages (Irvine and Schell, 2001; Shears, 2015; Livermore *et al*., 2016). InsP_6_, also known as phytate when chelated with metals, is the most abundant InsP species found in plants and is important for P*i* storage as well as signaling (Raboy *et al*., 2001). InsP_6_ is implicated as a structural component in plant auxin signaling through binding with the auxin receptor, transport inhibitor 1 (TIR1) (Tan *et al*., 2007). In plants, InsP_6_ can be acted on sequentially by two enzymes, the inositol 3,4,5,6-tetrakisphosphate 1-kinase (ITPK), and the diphosphoinositol pentakisphosphate 1-kinase (named VIP or VIH), to form the two known PP-InsPs, commonly referred to as InsP_7_ and InsP_8_ (Adepoju *et al*., 2019; Laha *et al*., 2019; Whitfield *et al*., 2020). InsP_7_ and InsP_8_ are hypothesized to bind and regulate the jasmonate receptor COI-1 (Laha *et al*., 2015; Laha *et al*., 2016) although InsP_5_ has also been implicated due to its presence within the crystallized COI-1 complex (Sheard *et al*., 2010). Understanding the specific roles of PP-InsPs in these and other plant signaling pathways is an area of emerging interest and requires further examination.

The role of PP-InsPs as critical players in eukaryotic P*i* sensing and homeostasis has been facilitated by examination of loss-of-function mutants in the InsP and PP-InsP synthesis pathways (Zhu *et al*., 2019; Dong *et al*., 2019; Land *et al*., 2021). Specifically, depleting InsP_8_ *in vivo* leads to alterations in a plant’s ability to properly sense and respond to P*i* (Zhu *et al*., 2019; Dong *et al*., 2019; Land *et al*., 2021). *Arabidopsis* loss-of-function mutants for both InsP_8_ synthetic enzymes VIP1 and VIP2 (Desai *et al*., 2014; Freed *et al*., 2020), which are also referred to as VIH2 and VIH1 (Zhu *et* al. 2019), have depleted levels of PP-InsPs, as well as altered physiological responses to P*i* which include increased P*i* accumulation, PSR gene induction, and impacted growth (Zhu *et al*., 2019; Dong *et al*., 2019). These data support a model for control of the PSR transcriptional response where InsP_8_ controls association of the Phosphate Starvation Response Regulator 1 (PHR1) transcription factor with its binding partner, SPX1, preventing PHR1-mediated induction of PSR genes. The SPX domain present within SPX1 has been shown via binding assays to bind InsP_8_ at a high affinity (Wild *et al*., 2016; Ried *et al*., 2021), thus suggesting InsP_8_ can be viewed as a proxy for intracellular P*i* levels and is likely the main molecule regulating PHR1-SPX complexes.

It is crucial to understand the extent to which InsPs and PP-InsPs contribute to plant P*i* sensing and to utilize this information to inform future translational approaches to circumvent a global P crisis. The overarching goal of our work is to develop a cover crop that can accumulate excess P*i* from polluted soils by leveraging PP-InsPs. A multitude of studies have demonstrated plants that can over accumulate P*i* by upregulating molecular components that are downstream of PP-InsP regulation, such as P*i* transporters (Rae *et al*., 2003; Jia *et al*., 2011; Zhang *et al*., 2014; Ye *et al*., 2015) and transcription factors (Yi *et al*., 2005; Nilsson *et al*., 2007; Matsui *et al*., 2013). Notably, a study by Rae *et al* demonstrates that increasing P*i* transporter expression does not always increase P*i* accumulation, as shown in their work where they overexpressed a high-affinity P*i* transporter AtPht1;1 in *Hordeum vulgare* L (barley) and did not observe a change in plant P*i* accumulation (Rae *et al*., 2004). Given this, we sought to establish a proof of concept of using decreased PP-InsPs to increase P*i* accumulation in *Arabidopsis* and in a cover crop species.

In developing a translational approach, we have assessed previously characterized *Arabidopsis* mutants with decreased PP-InsPs. While *vih1-2vih2-4* mutants could be considered since these mutants lack the ability to synthesize InsP_8_ (Zhu *et al*., 2019), the severe growth phenotypes of these mutants make it difficult to explore P*i* homeostasis in agronomic conditions. Instead, we have focused on developing a synthetic biology approach to directly target and break down PP-InsPs in plants. To achieve this, we used a yeast phosphatase to artificially remove plant PP-InsPs. We selected <u>d</u>iadenosine and <u>d</u>iphosphoinositol <u>p</u>olyphosphate phosphohydrolase (DDP1; gene YOR163w), a phosphatase from *Saccharomyces cerevisiae* containing a NUDIX hydrolase motif that hydrolyzes phosphoanhydride bonds from InsP_7_ and InsP_8_, polyphosphates (polyP), and diadenosine polyphosphates (Ap_n_A; specifically, Ap_5_A and Ap_6_A) (Safrany *et al*., 1999; Lonetti *et al*., 2011; Kilari *et al*., 2013; Márquez-Moñino *et al*., 2021). Of these three molecule classes, only PP-InsPs have been detected in plants (Desai *et al*., 2014; Laha *et al*., 2015) and have a known synthesis pathway (Desai *et al*., 2014; Laha *et al*., 2015; Adepoju *et al*., 2019; Laha *et al*., 2019).

We report that DDP1 overexpression in *Arabidopsis* decreases PP-InsP accumulation, increases P*i* accumulation over the course of plant development, and upregulates PSR gene expression. The physiological and developmental changes that occur in DDP1 transgenic plants are directly dependent on the level of DDP1 expression and the amount of P*i* present in the plant growth media. Further, we also transferred the *DDP1* gene into a cover crop species, *Thlaspi arvense* (pennycress), and show that this cover crop model system can increase P*i* accumulation from soil. Ultimately, these unique DDP1 *Arabidopsis* and pennycress transgenics will be useful in understanding the role of PP-InsPs in P*i* sensing as well as in other signal transduction pathways.

## RESULTS

### DDP1 overexpression in *Arabidopsis* negatively impacts growth

To reduce PP-InsPs *in planta*, transgenic lines were generated by overexpressing the *S. cerevisiae DDP1* gene (YOR163w) (Cartwright and McLennan, 1999; Lonetti *et al*., 2011; Kilari *et al*., 2013) fused to a C-terminal GFP tag under control of the CaMV 35S promoter in *Arabidopsis* Col-0 plants. DDP1 overexpression lines (DDP1 OX) were characterized for DDP1-GFP fusion protein expression by protein blotting with an anti-GFP antibody, which revealed correlation of DDP1- GFP protein of the expected molecular mass with a severe growth phenotype (Figure 1). For this study, we selected three DDP1 OX lines expressing abundant or very low amounts of DDP1-GFP. DDP1-A and DDP1-I transgenics have abundant DDP1-GFP expression and are severely impacted in their growth (Figure 1). These plants have a significantly reduced rosette diameter, as well as leaf chlorosis and leaf tip necrosis, which is evocative of previously characterized P*i* toxicity symptoms (Figure 1a-d) (Aung *et al*., 2006; Hu *et al*., 2011). As DDP1-I and DDP1-A plants begin the reproductive phase, abortion of developing embryos can be seen (Figure 1c), leading to low recovery of seeds. In contrast, the DDP1-H line has barely detectable levels of DDP1-GFP protein accumulation (Figure 1e), does not differ in growth from WT plants (Figure 1a, c), and does not have negatively impacted reproductive phenotypes. We compared growth of DDP1 OX lines with a commonly characterized PP-InsP-depleted mutant, *ipk1*. *ipk1* is a partial loss-of-function mutant for a gene encoding inositol pentakisphosphate 2-kinase, the only known enzyme in plants to synthesize InsP_6_ from InsP_5_ (Stevenson-Paulik *et al*., 2005). *ipk1* mutants are significantly smaller than WT plants, as seen by a reduced rosette diameter, which is a phenotype observed in past studies (Stevenson-Paulik *et al*., 2005; Kuo *et al*., 2014; Kuo *et al*., 2018). DDP1-I, DDP1-A, and to a lesser extent *ipk1* plants exhibit leaf tip chlorosis (Figure 1a-c), a phenotype previously linked to increased P*i* accumulation (Aung *et al*., 2006), and is suggestive of P*i* toxicity.

**Figure 1.**
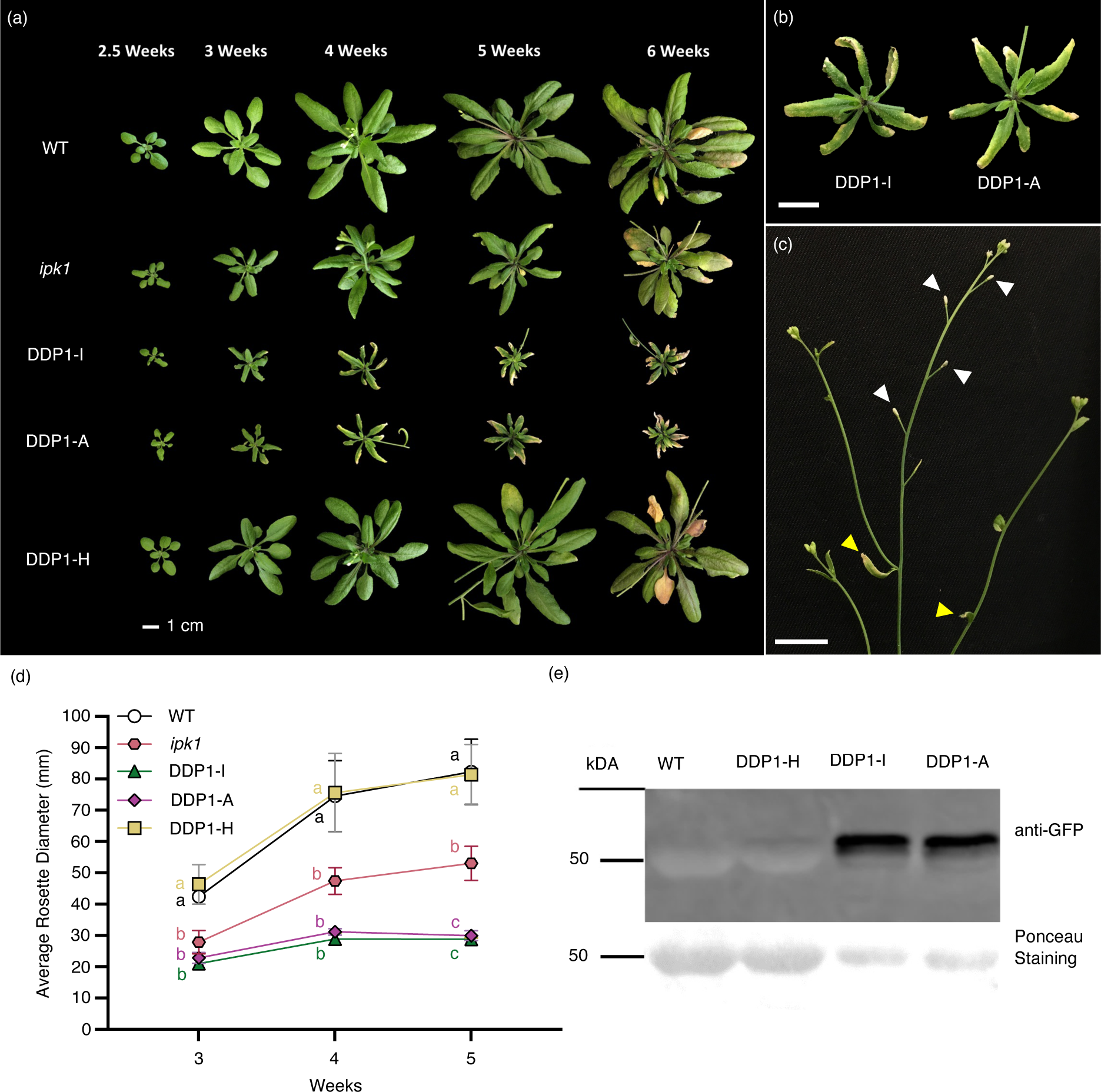
Characterization and Comparison of DDP1 OX Transgenics. (a) *Arabidopsis* rosette growth over the course of 6 weeks, each image representative of n = 3-4 independent experiments containing 12 or more plants per genotype. (b) Close up view of DDP1-I and DDP1-A rosettes and leaf tip necrosis and chlorosis after 4 weeks of growth. (c) Close up view of aborting siliques in 5- week-old in the DDP1-I line. White arrowheads indicate aborting siliques, yellow arrowheads mark yellowing cauline leaves. All Scale bars in (a-c) = 1 cm. (d) Average rosette diameter over time. Each point represents n = 3 independent experiments with over 12 plants per genotype per experiment; error bars show standard deviation (SD). Different letters indicate statistically significant means (Tukey HSD, α = 0.05). (e) Western blot of 4-week-old leaf tissue from WT and selected DDP1 OX transgenics. Ponceau staining shows RuBisCo accumulation in all plants as a positive control (∼50-56 kDa). WT and DDP1-H protein extracts were intentionally loaded to have double that of DDP1-I and DDP1-A to show the difference in DDP1-GFP accumulation, as shown by the Ponceau staining.

### DDP1 overexpression reduces seedling PP-InsP levels

To determine whether DDP1 overexpression alters PP-InsP profiles *in planta*, we measured InsPs and PP-InsPs in all three DDP1 OX lines and WT plants using our previously described *in vivo myo*-[^3^H] inositol radiolabeling method followed by high-performance liquid chromatography (HPLC) analysis (Desai *et al*., 2014; Adepoju *et al*., 2019). We observed a large decrease in InsP_7_ and InsP_8_ in both DDP1-I and DDP1-A plants, whereas DDP1-H profiles were similar to those of WT (Figures 2a and S1a). We compared the average percent of InsP_3_-InsP_8_ species in DDP1 OX transgenic lines to WT (Table 1). Looking at the consistent changes that occur in the severe lines (DDP1-I and DDP1-A), we see that InsP_7_ and InsP_8_ are the only InsP species that decreased reproducibly across all replicates compared to WT. DDP1-I and DDP1-A respectively had 53 ± 9% and 25 ± 10% InsP_7_ of the total WT InsP_7_ pool and 51 ± 4 % and 26 ± 11 % of the total WT InsP_8_ pool. Additionally, we observed an increase in the total InsP_4_ in both DDP1-I (144% ± 29%) and DDP1-A (157% ± 53%). We also expressed changes in PP-InsPs as ratios to make our data more easily comparable to that of other groups (Figures 2b-c, S1b) (Zhu *et al*., 2019). DDP1-A InsP_7_/InsP_6_ and InsP_8_/InsP_6_ ratios were significantly lower than WT ratios (Figure 2b-c). DDP1-I also had a significantly lower InsP_8_/InsP_6_ ratio compared to WT, though the InsP_7_/InsP_6_ ratio, while substantially lower compared to that of WT, was not significantly different. Interestingly, there was no significant change in the InsP_7_/InsP_8_ ratio in the severe DDP1 OX lines compared to WT, suggesting that the decrease in InsP_7_ and InsP_8_ are proportional (Figure S1b). Taken together, the data from growth analyses, protein expression, and PP-InsP profiling indicate that DDP1 OX lines with higher DDP1-GFP protein accumulation have large decreases in InsP_7_ and InsP_8_ and a small increase in total InsP_4_, which correlates with the observed growth phenotypes of reduced plant size and necrosis on leaf margins.

**Figure 2.**
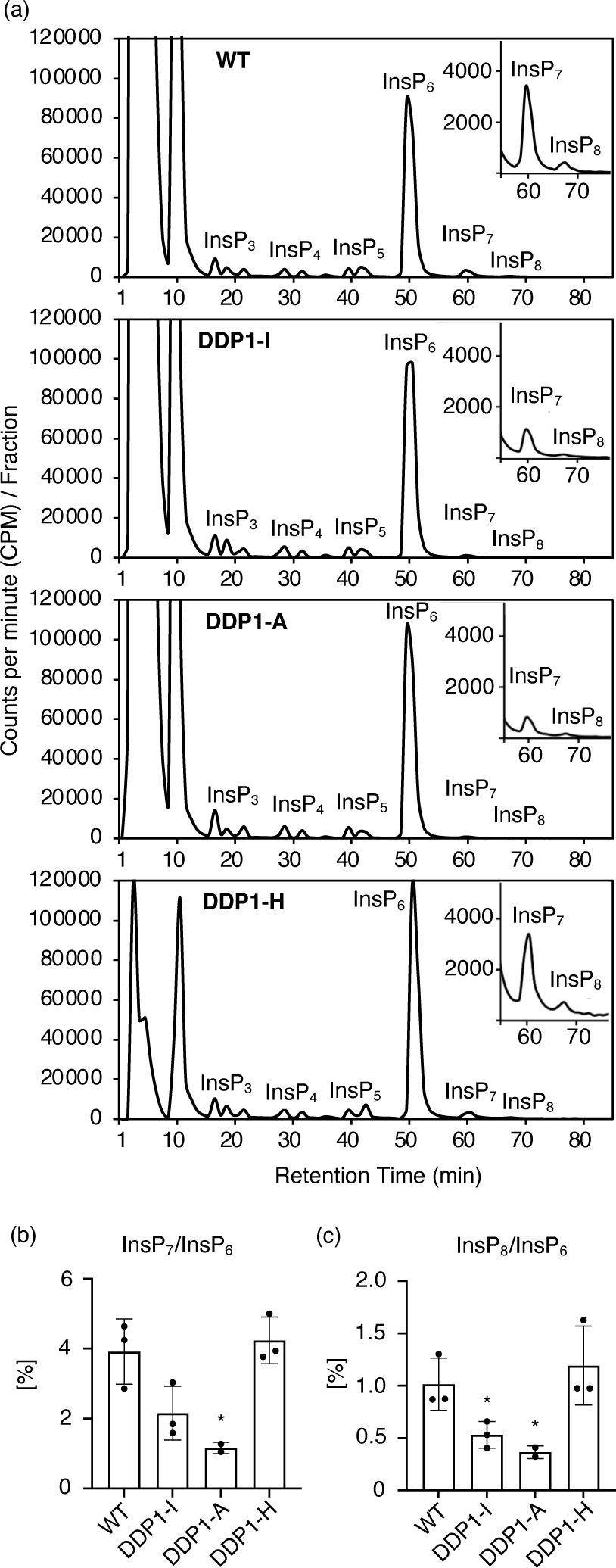
InsP and PP-InsP Profiling of DDP1 OX Transgenics. (a) WT and DDP1 OX transgenics were grown for 14 days on semi-solid 1/2x MS media with 0.2% agar then 100 μCi [^3^H]-*myo-*inositol was added for 4 days. All InsPs were extracted, separated using anion exchange HPLC, and data analyzed, as described in Experimental Procedures. These InsP profiles are representative of 2-3 independent replicates per genotype; see Figure S1 for all profiles. (b) InsP_7_/InsP_6_ and (c) InsP_8_/InP_6_ ratios. Asterisks show significant differences from WT; analyzed using Student’s *t*-test; * P < 0.05, error bars show SD of n = 2-3.

**Table 1.**
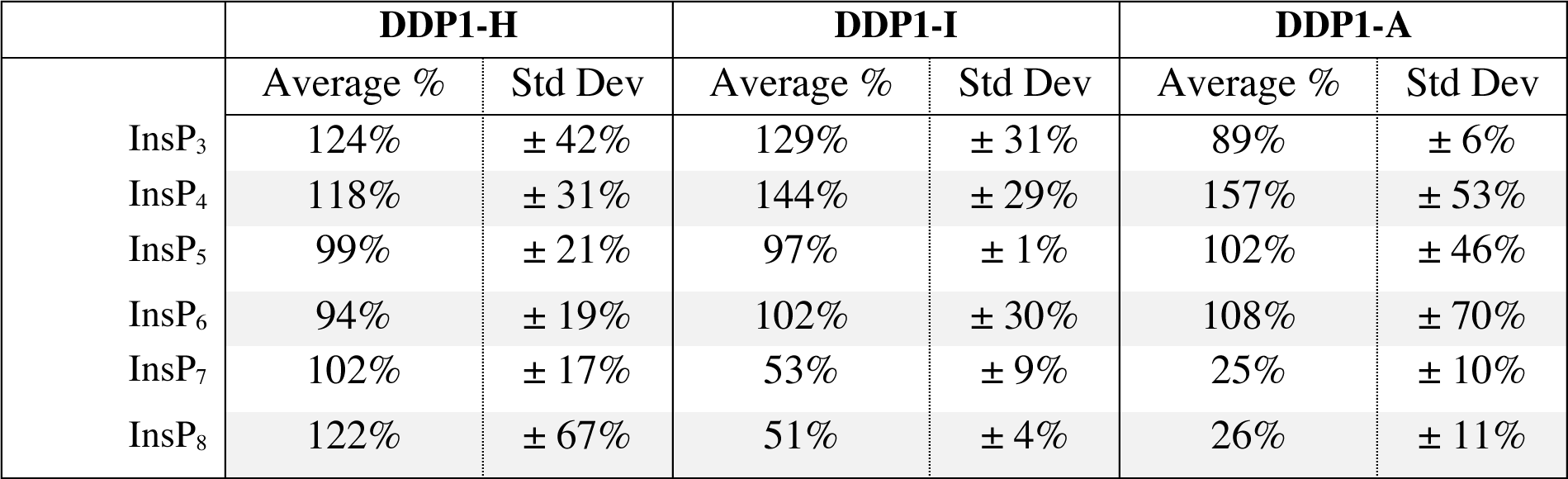
Average percent of the total CPM for each InsP as a percentage compared to the total WT pool for the respective InsP species. Averages are of all replicates shown in Figures 2a and S1a.

|  | **DDP1-H** |  | **DDP1-I** |  | **DDP1-A** |  |
| --- | --- | --- | --- | --- | --- | --- |
|  | Average % | Std Dev | Average % | Std Dev | Average % | Std Dev |
| InsP<sub>3</sub> | 124% | ± 42% | 129% | ± 31% | 89% | ± 6% |
| InsP<sub>4</sub> | 118% | ± 31% | 144% | ± 29% | 157% | ± 53% |
| InsP<sub>5</sub> | 99% | ± 21% | 97% | ± 1% | 102% | ± 46% |
| InsP<sub>6</sub> | 94% | ± 19% | 102% | ± 30% | 108% | ± 70% |
| InsP<sub>7</sub> | 102% | ± 17% | 53% | ± 9% | 25% | ± 10% |
| InsP<sub>8</sub> | 122% | ± 67% | 51% | ± 4% | 26% | ± 11% |

### DDP1-GFP localizes to the same cellular compartments as PP-InsP synthetic enzymes

Given the decrease in PP-InsPs in DDP1-I and DDP1-A but not DDP1-H, we were interested in whether DDP1 accumulated in the cellular compartments in which PP-InsPs putatively localize. While the exact cellular compartmentalization of PP-InsPs is unknown, multiple studies demonstrate that InsP and PP-InsP localize to the nuclei and cytoplasm (Xia *et al*., 2003; Kuo *et al*., 2018; Adepoju *et al*., 2019). Thus, we assess whether the subcellular localization of DDP1 in our plants was similar to enzymes in the PP-InsP synthesis pathway. Natively in yeast, DDP1 localizes to nuclei (Huh *et al*., 2003). However, it was important to establish this *in planta*. To assess DDP1 subcellular compartmentalization, we generated constructs expressing DDP1-GFP under control of the 35S CaMV promoter and assessed expression in *Arabidopsis* and *Nicotiana benthamiana*. We queried expression in two plant systems because we wanted to determine where DDP1-GFP was stably expressed in our *Arabidopsis* transgenics as well as in *N. benthamiana* to enhance localization resolution, co-localize with other proteins, and assess transient expression over the course of time.

In *Arabidopsis* transgenics, DDP1-GFP localizes to the nuclei and cytoplasm in leaf and root cells of DDP1-I and DDP1-A plants (Figures 3a-c and S2). DDP1-GFP signal is abundant in these lines, while DDP1-H samples had nearly undetectable DDP1-GFP signal (Figure S2d). Similarly in *N. benthamiana* leaves, DDP1-GFP transient expression results in accumulation of DDP1-GFP within the nucleus and cytoplasm of epidermal cells (Figures 3d-e, g-l, and S3). The pattern of nuclear and cytosolic localization is consistent 24-48 hours post-infiltration, suggesting that DDP1 could be breaking down PP-InsPs in the nucleus and/or cytoplasm. To ensure that this was an inherent property of DDP1 and not the position of the C-terminal GFP tag alone, we analyzed transient expression of DDP1 containing an N-terminal YFP tag in *N. benthamiana*. YFP- DDP1 localized to the same compartments as DDP1-GFP, indicating that DDP1 localizes to the epidermal cell cytoplasm and nuclei regardless of the location of the tag within the fusion construct (Figures 3f and S4). To increase confidence in cytosolic localization patterns, we co-expressed DDP1-GFP and YFP-DDP1 with two mCherry constructs: soluble mCherry, which localizes to the cytoplasm and nucleus, and ER-mCherry, a construct that localizes to the endoplasmic reticulum (ER) (Nelson *et al*., 2007). Using these in *N. benthamiana* allows one to distinguish between the ER and the cytoplasm as the ER can sometimes be closely associated with the nuclear membrane. Co-localization experiments show that DDP1-GFP and YFP-DDP1 colocalize with soluble mCherry in the nucleus and the cytoplasm and do not colocalize with the ER-mCherry tag (Figures 3g-l and S4-5). Collectively, our data demonstrates that DDP1 is localized to the cytoplasm and nucleus in both stable *Arabidopsis* cells and *N. benthamiana* cells.

**Figure 3.**
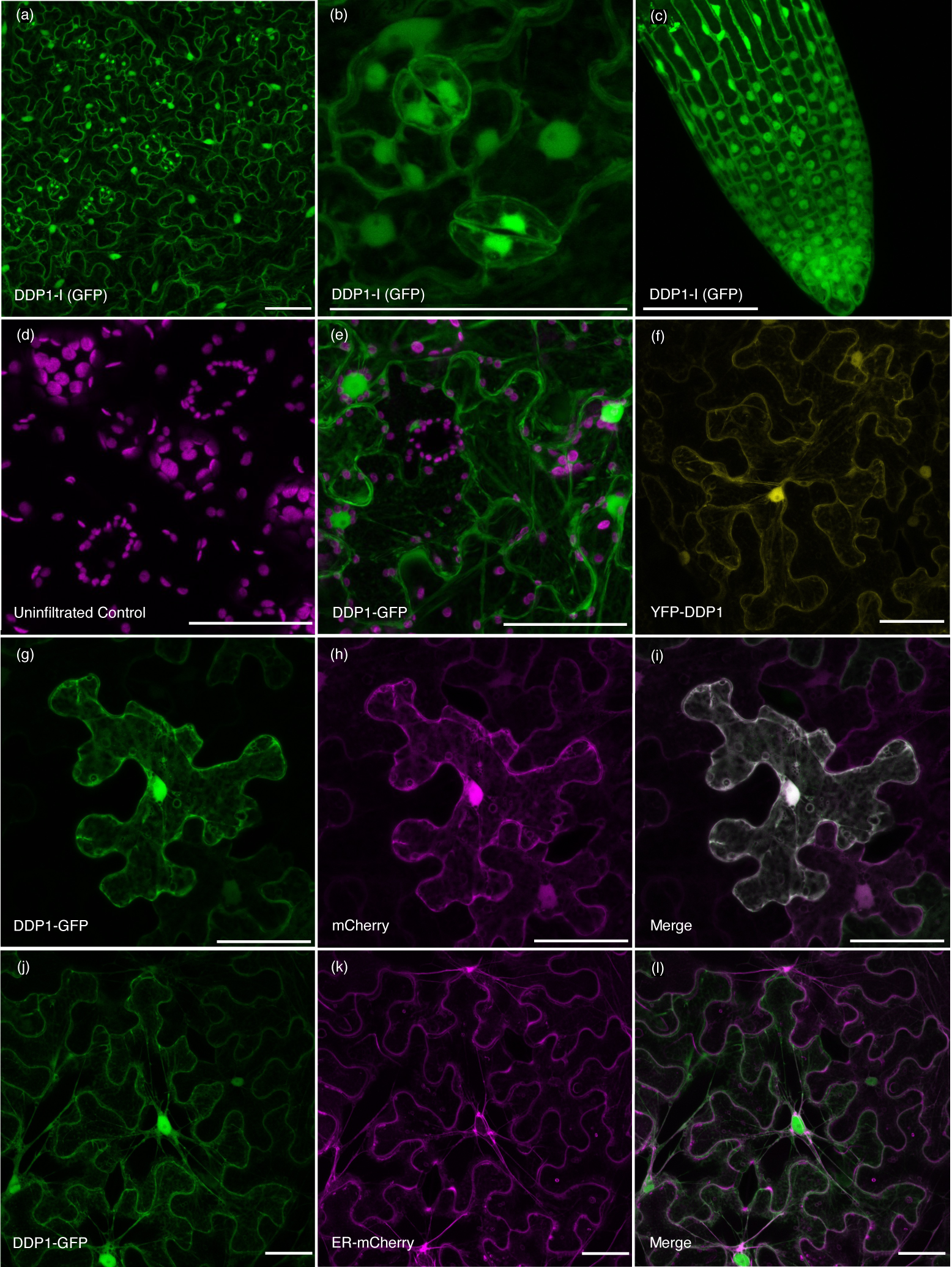
Confocal imaging of DDP1-GFP and YFP-DDP1 expression in Arabidopsis and *N. benthamiana*. DDP1-I leaves (a-b) and roots (c). (a) shows the palisade mesophyll cells and (b) magnifies the smaller guard cells. Uninfiltrated negative control *N. benthamiana* leaf (d) and DDP1-GFP transient expression in *N. benthamiana* leaves (e) 48 hours post infiltration. DDP1- GFP is shown in green and autofluorescence is shown in magenta. Transient expression of YFP- DDP1 (YFP channel; yellow) in *N. benthamiana* leaves 48 hours post-infiltration (f). *N. benthamiana* leaves co-infiltrated with DDP1-GFP and mCherry (g-i) or DDP1-GFP and ER- mCherry (j-l). All images are presented as maximum intensity projections from confocal Z-stack optical sections. All scale bars = 50 µm.

**Figure 4.**
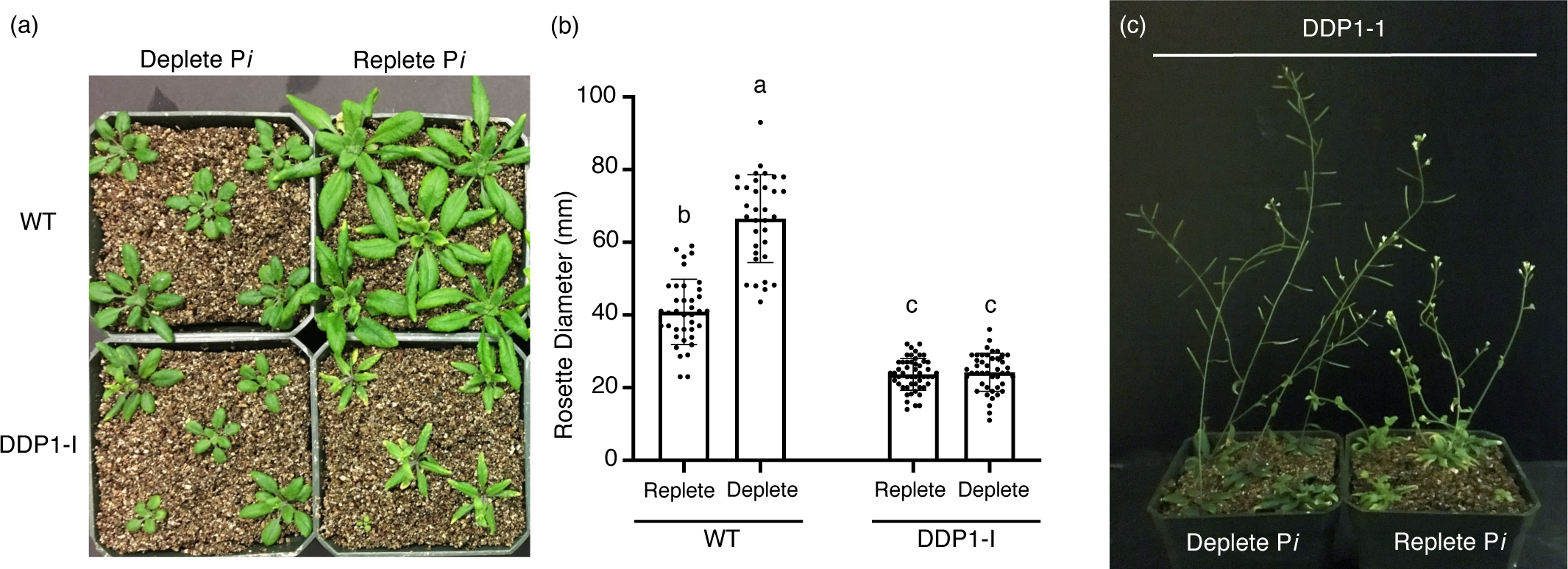
DDP1-I phenotypes grown under varying P*i* conditions. (a) WT and DDP1 OX transgenics grown on vermiculite containing 1/2x MS media with 10μM KH_2_PO_4_ (deplete P*i*) or 1 mM KH_2_PO_4_ (replete P*i*) after 35 days. DDP1-I plants did not accumulate lesions under deplete P*i* and resemble WT plants on deplete P*i*. (b) Rosette diameter measurements of WT and DDP1-I grown under deplete and replete P*i*. Each point represents an individual plant measurement from n = 2 independent experiments; over 35 plants per genotype and condition; error bars show SD. Different letters indicate statistically significant means (Tukey HSD, α = 0.05). (c) DDP1-I transgenics after 50 days of growth on vermiculite. DDP1-I lines grown on deplete P*i* did not have aborted siliques.

### DDP1 plants exhibit sensitivity to P*i*

Given the connection between PP-InsPs and P*i* sensing, we queried DDP1 OX P*i*-related phenotypes. To test how DDP1 OX plants respond to P*i*-deplete conditions, we focused on the DDP1-I line and grew plants using vermiculite supplemented with defined nutrient solutions containing either 10 μM or 1 mM KH_2_PO_4_ for up to 50 days to simulate P*i*-deplete and replete conditions, respectively. As expected, growth of WT plants is greatly impacted by P*i*-deplete conditions, with a significant reduction in shoot growth (Figure 4a-b). In contrast, we found that DDP1 OX transgenics did not show a significant difference in rosette diameter growth in P*i*- deplete as compared to P*i*-replete conditions though both were significantly smaller than WT under both conditions (Figure 4a-b). Additionally, while DDP1-I leaves are lighter in color and exhibit leaf tip necrosis and chlorosis in P*i*-replete conditions, growth under P*i*-deplete conditions results in improvement in growth as seen by a darker leaf color and absence of leaf tip necrosis and chlorosis (Figure 4a). A qualitative assessment of seed production indicated that growth of DDP1- I plants under P*i*-deplete conditions resulted in a rescue of the embryo abortive phenotype, as seen by DDP1 OX plants with stems and siliques that resembled WT plants after 50 days of growth (Figure 4c).

### DDP1 overexpression impacts P*i* accumulation and PSR gene expression

Previous studies show that perturbing PP-InsPs can increase P*i* accumulation, which is associated with an upregulation of PSR genes (Kuo *et al*., 2014; Kuo *et al*., 2018; Zhu *et al*., 2019; Laha *et al*., 2019; Land *et al*., 2021). As DDP1 OX phenotypes are P*i*-dependent and are similar to plants exhibiting symptoms of P*i* toxicity, we queried P*i* accumulation and expression of a specific set of PSR genes. We first gauged P*i* accumulation in soil grown DDP1 OX transgenics by measuring total P*i* accumulation from 3 to 7 weeks of growth in pooled leaf tissue (Figure 5a). Both severe DDP1 OX lines had an average fold increase of 5.5-6 times more P*i* compared to WT after 3 weeks of growth and, most extremely, a 25- to 26-fold-increase after 7 weeks of growth. This increase in P*i* accumulation in severe DDP1 OX lines was also significantly higher than P*i* accumulation in *ipk1* mutants (Figure 5a). This is notable as past studies demonstrate *ipk1* mutants accumulate significantly higher P*i* compared to WT (Stevenson-Paulik *et al*., 2005; Kuo *et al*., 2014; Kuo *et al*., 2018; Land *et al*., 2021). Under our conditions, *ipk1* mutants had higher P*i* from WT only after 4 weeks of growth whereas all other *ipk1* timepoints showed no statistical difference in P*i* accumulation (Figure 5a). WT-like DDP1-H transgenics also showed no difference in P*i* accumulation compared to WT plants. Most strikingly, a comparison of severe DDP1 OX transgenic and *ipk1* P*i* accumulation over time indicates that while *ipk1* plants reach a maximum at 3 weeks, DDP1 OX plants continue to accumulate significantly elevated levels of P*i* over time as compared to WT plants.

**Figure 5.**
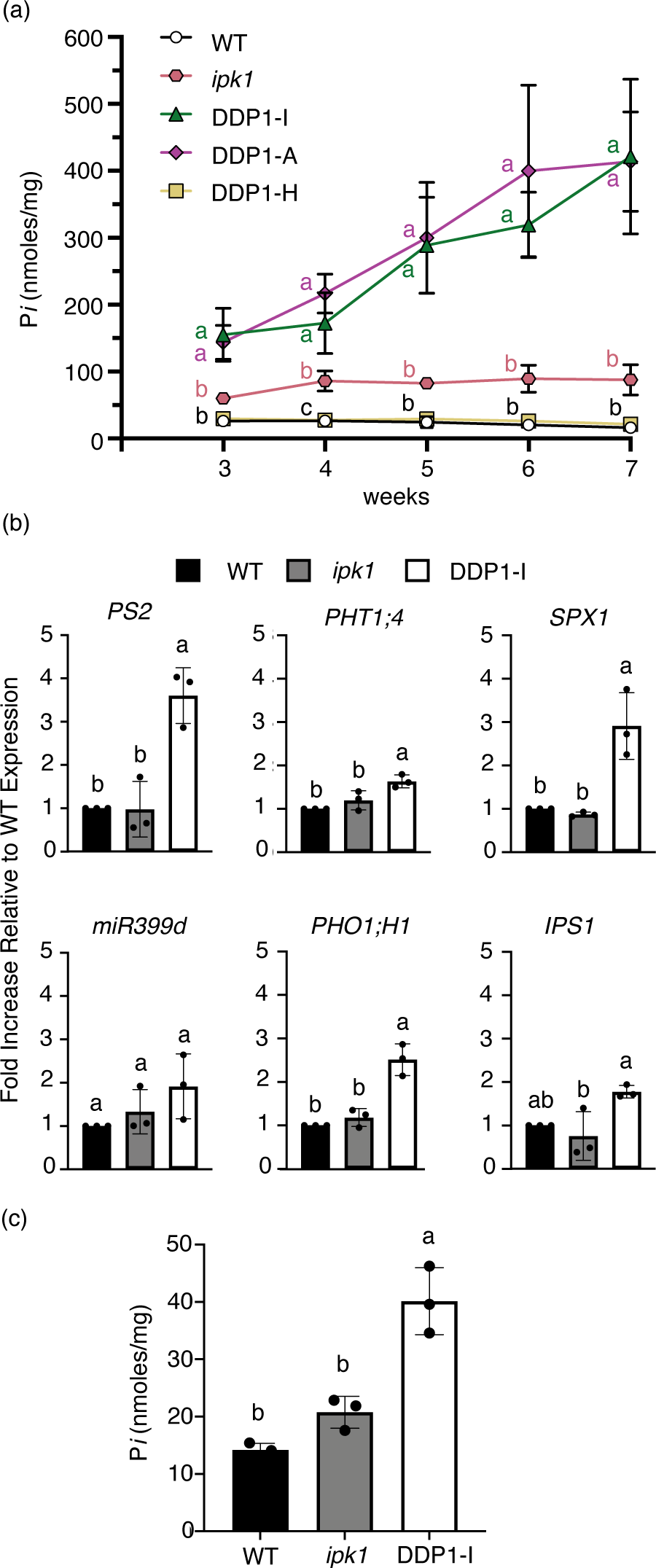
P*i* accumulation and PSR gene induction in DDP1 OX lines. (a) P*i* accumulation in pooled leaf tissue from 3-7-week-old soil-grown plants. Each point represents pooled plant tissue from n = 2-4 independent experiments; error bars show SD. Different letters indicate statistically significant means (Tukey HSD, α = 0.05). (b) PSR gene expression (relative to WT) and (c) P*i* accumulation in whole seedlings grown for 10 days grown on media plates. Error bars denote SD of n = 3 independent experiments. Different letters indicate statistically significant means (Tukey HSD, α = 0.05).

We hypothesized that increased P*i* accumulation over time was tied to increased expression of PSR genes caused by a reduction of PP-InsPs. We queried a set of commonly used PSR marker genes that play key roles in plant P*i* accumulation under P*i*-deplete conditions (*PS2, PHT1;4*, *SPX1*, *miR399d*, *PHO1;H1*, *IPS1*) (Rouached *et al*., 2010; Jost *et al*., 2015; Chien *et al*., 2018). Of the genes queried, we observed that 10-day-old DDP1 OX whole seedlings showed a significant upregulation of *PS2*, *PHT1;4*, *SPX1*, and *PHO1;H1* compared to WT and *ipk1* seedlings (Figure 5b). DDP1-I also had significantly increased *IPS1* compared to *ipk1* however this change was not significantly different from WT. *ipk1* seedlings trended towards increased expression of a few genes within the queried subset, however these changes were not statistically significant. This result was expected as *ipk1* whole seedlings have not been shown to induce the PSR under replete conditions (Zhu *et al*., 2019), though it is important to consider that previous studies querying PSR gene expression specifically in *ipk1* roots show significant increases in PSR gene expression (Kuo *et al*., 2014; Kuo *et al*., 2018). P*i* accumulation in these 10-day-old seedlings was also queried and we found, as expected, that DDP1-I plants accumulated significantly more P*i* compared to WT and *ipk1* (Figure 5c).

### DDP1 overexpression in pennycress yields P*i*-accumulation phenotypes

Based on the unique ability of DDP1 OX plants to accumulate higher amounts of P*i* over time, we were interested in translating this discovery into a crop species to accumulate excess P*i* from soil as a potential phytoremediation utility. We selected *Thlaspi arvense* (pennycress), a winter annual cover crop that is commonly used by farmers and shows immense promise as a biofuel crop (Chopra *et al*., 2020). It is important to note that great efforts have been made to establish pennycress as a model cover crop system through its recently sequenced genome and assembled transcriptome, making it an ideal system for translation of our findings in *Arabidopsis* DDP1 OX transgenics (McCormick, 2018). We demonstrated that pennycress plants can be stably transformed to overexpress DDP1 and show similar phenotypes as *Arabidopsis* DDP1 OX transgenics (Figure 6). Notably, we characterized heterozygotes in this study as homozygous pennycress lines were too difficult to grow based on extreme P*i* toxicity (Figure S6). After 5 weeks of growth, the DDP1-B OX line with higher transgenic protein accumulation manifested leaf tip necrosis and chlorosis phenotypes (Figure 6a-c). Our pennycress DDP1 OX transgenics overexpress DDP1 with an N-terminal YFP tag, which accumulates in the same subcellular compartments (nuclear and cytosolic) as in *Arabidopsis* DDP1 OX plants and *N. benthamiana* leaf tissue (Figure 6d). Most prominently, pennycress DDP1-B OX plants exhibit significantly higher P*i* accumulation compared to WT and the WT-like DDP1-L OX line (Figure 6e). Taken together, these data confirm that overexpression of DDP1 in both *Arabidopsis* and pennycress share a similar mechanism through common DDP1 subcellular localization, manifestation of similar P*i* toxicity phenotypes, and enhanced P*i* accumulation in leaf tissue.

**Figure 6.**
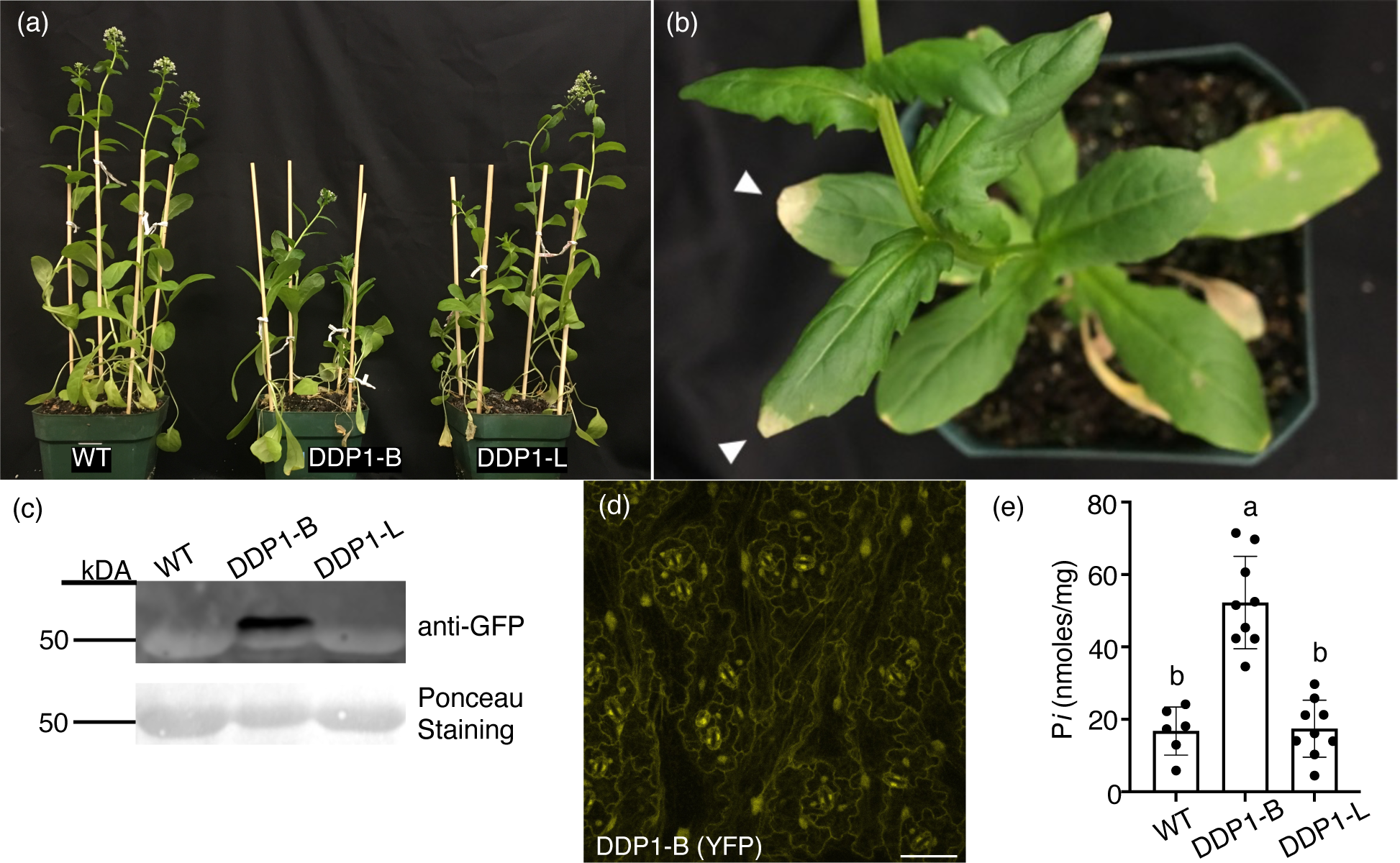
Characterization of Pennycress DDP1 OX Transgenics. (a) Physiology of WT pennycress and two independent, heterozygous DDP1 OX transgenic lines after 5 weeks of growth. (b) Close-up view of DDP1-B leaf lesions after 7 weeks of growth on soil. White arrowheads mark yellowing of leaf tips. (c) Western blot of 6-week-old leaf tissue from WT and heterozygous pennycress DDP1 OX lines. Ponceau staining shows RuBisCo accumulation in all plants as a positive control (∼50-56 kDa). (d) Stable YFP-DDP1 expression in pennycress DDP1-B leaves. Scale bar = 50 µm. (e) P*i* accumulation in 5.5 to 7-week-old pennycress leaf tissue. Error bars denote SD of n = 3 independent experiments; each point represents leaf tissue of an individual heterozygous plant. Different letters indicate statistically significant means (Tukey HSD, α = 0.05).

## DISCUSSION

PP-InsP signaling in plants has become an increased area of interest due to emerging evidence connecting the role of PP-InsPs to P*i* sensing and other hormone signaling pathways (Laha *et al*., 2015; Kuo *et al*., 2018; Adepoju *et al*., 2019; Laha *et al*., 2019; Zhu *et al*., 2019; Laha *et al*., 2020; Land *et al*., 2021). In this work, we sought to use a synthetic biology approach to manipulate PP-InsPs in transgenic plants to study the long-term consequences of decreased PP-InsPs, and to evaluate the preliminary translational potential of depleting PP-InsPs *in planta*. This work is novel as it artificially depletes PP-InsPs to leverage P*i* accumulation in two plant species. The two most notable findings from this study are that DDP1 overexpression decreases PP-InsPs *in planta* and confers a unique P*i*-accumulating phenotype with promising translational application. It is also important to note that our study queries the long-term mechanistic impacts of a deregulated plant PSR by measuring P*i* accumulation of soil-grown plants over the course of multiple weeks.

### DDP1 overexpression results in unique biological characteristics distinct from PP-InsP synthesis mutants

DDP1 overexpression *in planta* allowed us to achieve our original goal of removing PP-InsPs. This strategy is distinct from studying InsP and PP-InsP kinase loss-of-function mutants. While there are similarities in how PP-InsPs are altered in DDP1 OX plants and previously characterized PP-InsP synthesis mutants, DDP1 OX transgenics likely have actively hydrolyzed PP-InsPs in contrast to loss-of-function mutants that lack enzymes that synthesize InsP_6_ and/or PP-InsPs. There are three types of synthesis mutants with defects in P*i* sensing that have been described in current literature; the functions of the WT encoded enzyme products of these mutants are shown in Figure 7. The well-studied *ipk1* mutant used in this and other studies is not a complete loss-of-function mutant for IPK1, as complete *ipk1* nulls are nonviable (Stevenson-Paulik *et al*., 2005; Kuo *et al*., 2014). To this end, *ipk1* mutants synthesize greatly reduced levels of InsP_6_, resulting in reduced levels of PP-InsPs (Laha et al., 2015; Kuo *et al*., 2018; Land *et al*., 2021). Similarly, *itpk1* mutants showed a reduction in InsP_7_ (Kuo *et al*., 2018) and InsP_8_ (Laha *et al*., 2020) compared to WT seedlings. *vih1-2vih2-4* mutants were reported to have undetectable levels of InsP_8_ and double the WT pool of InsP_7_ (Zhu *et al*., 2019). This buildup of InsP_7_ is likely attributed to the fact that InsP_7_ cannot be converted to InsP_8_. All three types of characterized mutants showed P*i*-related growth phenotypes, increases in leaf P*i* accumulation compared to WT plants, and an induction of PSR genes (Stevenson-Paulik *et al*., 2005; Kuo *et al*., 2014; Kuo *et al*., 2018; Zhu *et al*., 2019). This induction of the PSR is thought to result from the lack of InsP_8_ available within these mutants, which acts to release the PHR1 transcription factor (Figure 7).

**Figure 7.**
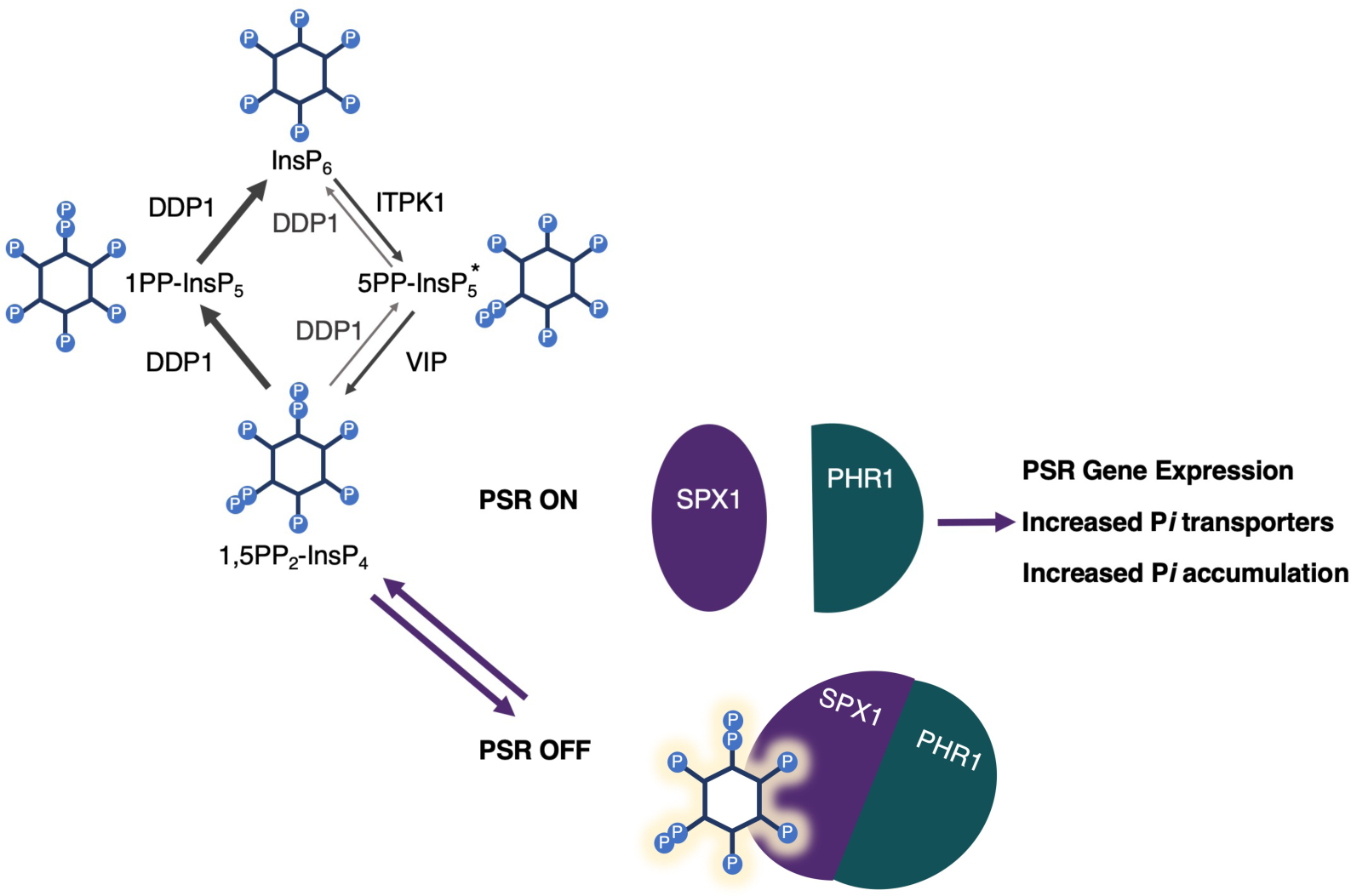
Model for cyclical degradation and synthesis of PP-InsPs in DDP1 transgenics and PSR gene regulation. DDP1 likely hydrolyzes 1,5PP_2_-InsP_4_ (InsP_8_) and 1-InsP_7_ *in Planta* as well as 5-InsP_7_ to a lesser extent (depicted in gray). The * depicts the plant predominant form of InsP_7_. Native plant ITPK1 and VIP enzymes catalyze synthesis of 5-InsP_7_ and 1,5 PP_2_-InsP_4_, though measurement of PP-InsPs in DDP1 plants suggests the rate of degradation exceeds the rate of synthesis. Decreased 1,5PP_2_-InsP_4_ accumulation results in dissociation of SPX–PHR complexes, enabling PHR1 to bind to PSR gene promoters and turn on the PSR. Not shown are the recently identified structural changes that suggest InsP_8_ binding to SPX1 controls the oligomeric state of PHR1 complexes (Ried et al., 2021).

Despite the strength of data implicating that InsP_8_ is the sole molecule regulating P*i* sensing (Zhu *et al*., 2019; Dong *et al*., 2019; Ried *et al*., 2021), not all reported genetic mutants fit in with this currently accepted model. One example of this is the *Arabidopsis vih2-4* mutant. It has been demonstrated that *vih2-4* mutants have lower InsP_8_ levels and higher InsP_7_ levels (Laha *et al*., 2015), but do not have changes P*i* accumulation or an induction in PSR genes (Kuo *et al*., 2018). It is also important to consider that *itpk4-1*, an *Arabidopsis* mutant with decreased InsP_6_, InsP_7_, and InsP_8_ levels, did not show a significant increase in P*i* accumulation nor induction of PSR genes (Kuo *et al*., 2018; Wang *et al*., 2021). Additionally, a recent study shows that a separate *Arabidopsis* VIP mutant, *vip1-2/2-2*, has 70-75% of the total WT InsP_8_ level and no significant changes in InsP_7_ (Land *et al*., 2021). Surprisingly, *vip1-2/2-2* seedlings accumulate significantly lower levels of P*i* and *vip1-2/2-2* leaves have no significant difference in P*i* accumulation compared to WT and *ipk1 mutants*. PSR genes in *vip1-2/2-2* mutants are also downregulated, rather than upregulated (Land *et al*., 2021). The lack of correlation between InsP_8_ and P*i* sensing in this broad set of mutants indicates the possibility that another InsP besides InsP_8_, or the ratio of InsPs/PP-InsPs, might play an important role in P*i* sensing.

Another key difference between our DDP1 OX plants and *ipk1* mutants is the level and duration of P*i* accumulation (Figure 5a). While our work and that of others shows that *ipk1* mutants accumulate higher P*i* compared to WT plants under specific conditions (Stevenson-Paulik *et al*., 2005; Kuo *et al*., 2014; Kuo *et al*., 2018; Land *et al*., 2021), expression of DDP1 in plants allows for a much greater, and continual accumulation of P*i* (Figure 5a). We note that this increased P*i* accumulation is associated with a greater induction of PSR genes in DDP1 as compared to *ipk1* seedlings (Figure 5b). It is important to note that previous work on *ipk1* mutants measured PSR gene expression in roots (Kuo *et al*., 2014; Kuo *et al*., 2018), while our work has focused on the whole seedling impact of altering PP-InsPs, as the leaf tissue present the greatest opportunity with regards to future phytoremediation strategies. In addition to our work, Zhu et al shows that *ipk1* whole seedlings do not induce a similar queried subset of PSR genes (Zhu *et al*., 2019). Thus, it seems likely that our DDP1 OX plants have a stronger induction of the PSR compared to *ipk1* plants.

### DDP1 functionality *in planta*

We show that stable, abundant expression of DDP1 in *Arabidopsis* consistently decreases InsP_7_ and InsP_8_ (Figures 1 and 2). This decrease in PP-InsPs was expected given previous experiments characterizing DDP1 enzymatic activity on InsP_7_ and InsP_8_ (Safrany *et al*., 1999; Kilari *et al*., 2013). Based on the synthetic pathway in plants, it is thought that the predominant form of InsP_8_ is 1,5-InsP_8_ (also known as 1,5-[PP]_2_-InsP_4_ or 1,5-bis-diphosphoinositol 2,3,4,6-tetrakisphosphate) (Zhu *et al*., 2019; Adepoju *et al*., 2019; Laha *et al*., 2019). InsP_7_ has two different enantiomers: 1- InsP_7_ (also known as 1-PP-InsP_5_; 1-diphospho-*myo*-inositol 2,3,4,5,6-pentakisphosphate) and 5- InsP_7_ (also known as 5-PP-InsP_5_; 5-diphospho-*myo*-inositol 1,2,3,4,6-pentakisphosphate). While data from earlier literature suggested that 1-InsP_7_ rather than 5-InsP_7_ is the ideal substrate for DDP1 (Lonetti *et al*., 2011), a more recent study shows that DDP1 can also hydrolyze 5-InsP_7_, and preferentially hydrolyzes the 5-position PP bond from 1,5-InsP_8_ rather than from the 1-position *in vitro* (Kilari *et al*., 2013). A very recent and extensive crystallography study provides a molecular explanation for this in that PP-InsPs have several binding modes in the DDP1 catalytic pocket, with some being more productive for hydrolysis than others (Márquez-Moñino *et al*., 2021). This work established that 1-InsP_7_ binds in a more productive mode, whereas 5-InsP_7_ molecules bind less productively within DDP1. Márquez-Moñino *et al* also propose that 1,5 -InsP_8_ likely binds in two different modes, one of which is more productive for hydrolysis of the 5-PP. Given this, and our result that severe DDP1 OX profiles contain lower InsP_7_ and InsP_8_ levels compared to WT (Figure 2), we propose the following model for DDP1 activity within our transgenics (Figure 7): 1. We hypothesize that DDP1 is breaking down 1,5-InsP_8_ and 1-InsP_7_ as well as 5-InsP_7_ to a lesser extent. The increase in total InsP_4_ could be a result of increased InsP_7_ and InsP_8_ degradation. 2. Synthesis driven by native ITPK1 and the VIP enzymes may be replenishing the InsP_7_ and InsP_8_ pools, allowing for viability of DDP1 transgenic plants. 3. 5-InsP_7_ most likely accounts for the remaining InsP_7_ levels in our HPLC profiles. This model also aligns with the original suggestion that PP-InsP turnover is a cyclical interconversion between InsP_6_, InsP_7_ enantiomers, and InsP_8_ based on current evidence in yeast and humans (Shears, 2018). Thus, we have likely tipped the delicate scale of this continual cycle of PP-InsP degradation and synthesis by adding a nonendogenous phosphatase to the mix.

One other key finding of DDP1 action in plants stems from our localization of DDP1 to the nucleus and cytoplasm. While the specific locations of PP-InsPs have yet to be determined, DDP1 shows the same localization patterns as other known InsP and PP-InsP synthetic enzymes (Xia *et al*., 2003; Kuo *et al*., 2018; Adepoju *et al*., 2019), and is consistent with the native DDP1 localization in yeast (Huh *et al*., 2003). Thus, the similarity in subcellular localization patterns described here strongly support the role of DDP1 in targeting cytosolic and nuclear pools of PP-InsPs in *Arabidopsis* and pennycress. Although DDP1 can also act on other substrate signaling molecules in yeast and animals, there is currently no evidence to suggest that these other potential substrates are present in plants. These alternative substrates include two distinct types of molecules: polyphosphates (polyP; polyP_n_) and diadenosine polyphosphates (Ap_n_As; specifically, Ap_5_A and Ap_6_A), which are known substrates for DDP1 in *S. cerevisiae* (Safrany *et al*., 1999; Lonetti *et al*., 2011; Andreeva *et al*., 2019; Márquez-Moñino *et al*., 2021). Both molecular classes have also been linked to maintaining P*i* metabolism and cellular homeostasis in a variety of prokaryotic and eukaryotic organisms (Lorenzo-Orts *et al*., 2020; Pietrowska-Borek *et al*., 2020). Two studies report careful efforts made to detect and quantify these molecules in higher plants, which were unsuccessful (Pietrowska-Borek *et al*., 2011; Zhu *et al*., 2020). Given this, we hypothesize that the most likely action of DDP1 when expressed in plants is hydrolysis of PP-InsPs leading to the observed impacts on P*i* homeostasis. If true, this would point to key differences between plants, yeast, and animals regarding these signaling molecules and the roles they play *in vivo*.

### From proof of concept to potential phytoremediation strategy?

It is essential that we develop unique strategies to reduce fertilizer inputs and watershed pollution to circumvent future P*i* shortages (Cordell *et al*., 2009). Our expression of DDP1 in both a model plant and cover crop provide unique germplasm to study how decreased PP-InsPs impact plant growth, physiology, and P*i* accumulation without lethality. Our work is unique as we synthetically modulate PP-InsPs to increase P*i* accumulation. While a multitude of studies demonstrate that molecular components downstream of PP-InsP regulation can be exploited to increase plant P*i* accumulation (Rae *et al*., 2003; Yi *et al*., 2005; Nilsson *et al*., 2007; Jia *et al*., 2011; Matsui *et al*., 2013; Zhang *et al*., 2014; Ye *et al*., 2015), overexpression of plant transporters does not always lead to P*i* accumulation (Rae *et al*., 2004), suggesting that it will be more impactful to target regulators of P*i* transporters and transcription factors. Other common strategies employ key players in the InsP pathway by targeting and limiting InsP_6_ content in seeds and grains, as non-ruminant livestock animals cannot digest InsP_6_ and so most is excreted as waste (Sharpley and Withers, 1994; Abelson, 1999; Raboy, 2002). Should we alter InsPs and PP-InsPs in crop species, we must also consider how decreased PP-InsPs will impact how a crop responds to pathogens given the demonstrated ability of InsP_7_ and InsP_8_ bind to the COI1 jasmonate receptor complex (Laha *et al*., 2015; Laha *et al*., 2016). One study demonstrates that decreasing InsP_8_ impacts transcription of genes involved in jasmonic acid signaling and these plants were more susceptible to insect herbivory (Laha *et al*., 2015). Further, plants with alterations in InsP_6_ and PP-InsPs are impacted in their response to pathogens through alterations in salicylic acid signaling. A separate study shows that *Arabidopsis* mutants and transgenic potato plants with decreased InsP_6_ levels have an increased susceptibility to pathogens (Murphy *et al*., 2008). However, a recent study from Laha *et al* shows that *ipk1*, *itpk1*, and *vih2* mutants have enhanced resistance to pathogen infection and increased salicylic acid levels (Gulabani *et al*., 2021). Taken together, future strategies leveraging PP-InsPs in crop plants will need to consider how alterations in PP-InsPs will impact herbivory and pathogen susceptibility.

The P*i* crisis is a complicated issue that will require a variety of innovative strategies to solve. As the global population increases, there is a need for farmers to increase fertilizer inputs as crop production also increases. Between global food demand and higher fertilizer usage in agricultural and urban areas, there is a subsequent increase in fertilizer runoff into the watersheds. As we continue to learn more about the roles of PP-InsPs in plant P*i* sensing and growth, this will generate knowledge for developing unique strategies, ranging from P*i* removal from polluted environments to improving P*i*-use efficiency. While the growth tradeoff we characterized is not desirable, our proof of principle approach using DDP1 transgenic plants establishes a first step in altering PP-InsPs *in planta* leading to continual accumulation of P*i* from soil. This is important for the future development of unique plants to remediate P-polluted environments.

## EXPERIMENTAL PROCEDURES

### Cloning and Transformation

*DDP1* (YOR163W) from Saccharomyces cerevisiae was PCR amplified for ligation into Gateway pENTR/D-TOPO destination vector. Using the Gateway® LR Clonase™ II kit (Invitrogen Corp., Carlsbad, CA), *DDP1* was recombined with Gateway plant destination vectors pK7FWG2 (C-terminus DDP1-GFP) or pH7YWG2.0 (N-terminus YFP-DDP1) (VIB UGent Center for Plant Systems Biology). *E. coli* transformed with these constructs was selected on antibiotic media. Plasmids purified from resulting colonies were sequencing for verification of correct cloning. *Agrobacterium* (strain GV3101) was transformed with these constructs and were used for transient *Nicotiana benthamiana* experiments or to transform *Arabidopsis thaliana* (ecotype Col-0) plants. Transgenic *Thlaspi arvense* (Pennycress Spring 32-10; CS29065) were generated as described (McGinn *et al*., 2019), using the pHY7YWG2.0 N-terminus YFP-DDP1.

### Plant growth conditions

*ipk1* (SALK_065337; AT5G42810) T-DNA mutants were obtained from the Arabidopsis Biological Resource Center (ABRC, Columbus, OH, USA). All mutants, transgenics, and WT *Arabidopsis* lines were identically grown in concert on soil under long day conditions (16 hours light/8 hours dark, 55% humidity, day/night temperature 23/21°C and 120 μE light intensity).

Pennycress plants (Spring32-10; CS29085) were obtained from ABRC and grown under the same conditions as *Arabidopsis* transgenics with 300 μE light intensity. Pennycress transgenics were selected using hygromycin as they are naturally kanamycin resistant. Seedlings grown under sterile conditions were first sterilized using 100% ethanol for one minute, transferred to a 30% (v/v) Clorox solution for 5 minutes and washed five times with ddH_2_O. Seeds were placed in 0.5X MS media + 0.2% agar and stratified for 3 days at 4°C. Seeds were either transferred to 0.5X MS solid media plates or multi-well plates containing semi-solid media (0.2% agar). For the hydroponics experiment, stratified seeds were added to a solution containing 0.01% agarose and placed on top of pots containing washed vermiculite. All pots were placed in trays containing 2 liters of 0.25X MS media and varying P*i* (deplete (10 μM) and replete (1mM) KH_2_PO_4_). The rosette diameters were measured in all plants grown on soil and hydroponics. All data was analyzed using JMP Pro and GraphPad Prism.

### Protein Blots of GFP-Fusion Proteins

Protein blots were performed as previously reported (Burnette *et al*., 2003). Leaf tissue from soil grown plants was pulverized in liquid nitrogen, homogenized, and separated from cell debris. SDS- bromophenol blue loading dye was added to the extracted proteins and boiled for 5 minutes at 85°C. After a subsequent centrifugation, the supernatant was loaded onto a polyacrylamide gel; equal amounts of protein were added from each sample. For western blotting, a 1: 5000 dilution of anti-GFP antibody (Invitrogen Molecular Probes, Eugene, OR) and a 1: 2000 dilution of secondary goat anti-rabbit horseradish peroxidase antibody (Bio-Rad Laboratories, Hercules, CA) were used to detect GFP.

### InsP and PP-InsP Profiles of *Arabidopsis* Seedlings

WT and DDP1 OX transgenics were grown in semi-solid 0.5X MS media and 0.2% agar for two weeks, 30 μE light intensity. 24 seedlings of each genotype were transferred to Eppendorf tubes containing 300 μL of 0.5X MS media, 0.2% agar and 100 μCi of [^3^H] *myo*-inositol (20 Ci/mmol; American Radiolabeled Chemicals (ARC), St. Louis, MO, USA) for 4 days. InsPs were extracted from seedlings by pulverizing tissue in extraction buffer (1 M Perchloric acid (HClO_4_), 3 mM EDTA, and 0.1 mg/ml InsP_6_) and vortexed with glass beads. The pH of the extract was neutralized to pH 6-8 using a neutralization buffer (3 mM EDTA and 1M K_2_CO_3_). Samples were dried to a volume of 80-100 μL. Negatively charged species were separated via HPLC using a 125 x 4.6 mm Partisphere SAX-column (Sepax Technologies, Delaware, USA) and an ammonium phosphate elution gradient (Azevedo and Saiardi, 2006). Radioactivity in all collected fractions was quantified using a scintillation counter.

### Subcellular Localization and Imaging

All *Arabidopsis*, pennycress, and *N. benthamiana* cells were imaged using a Zeiss LSM 880 confocal microscope (Carl Zeiss) equipped with a 25x C-Apochromat water immersion lens. *N. benthamiana* plant leaves were infiltrated with transformed with *Agrobacterium tumefaciens*, as previously described (Kapila *et al*., 1997). Specifically, *Agrobacterium* cells were grown overnight, pelleted, and resuspended in MMA (10 mM MES, 10 mM MgCl_2_, 200 µM acetosyringone) solution at an OD_600_ of 1.0. After a 2-4 hour incubation period in the dark, *N. benthamiana* leaves were infiltrated with the *Agrobacterium* MMA cultures. Leaf sections were imaged after 24, 48, and 72-hours post infiltration. GFP was excited using a 488-nm argon laser and its fluorescence was detected at 500- to 550-nm and YFP was excited using a 488-nm argon laser and its fluorescence was detected at 517- to 561-nm. GFP- and YFP-tagged proteins were colocalized with a set of mCherry tagged organelle markers (Nelson *et al*., 2007), mCherry was imaged using excitation with a 594-nm laser and fluorescence was detected at 600- to 650-nm. Chlorophyll autofluorescence was excited using a 594-nm laser and emission above 650 nm was collected.

### P*i* accumulation assays

P*i* was extracted from ∼50 mg from *Arabidopsis* and pennycress tissue of various ages. Samples were pulverized in liquid nitrogen. A 1:10 ratio of 1% acetic acid was added to each sample, which was vortexed and incubated on ice, and centrifuged. Assays were performed on P*i* extracts using a modified micro-titer assay as previously reported (Ames, 1966). 50 μL of the supernatant and 1 mL of working reagent (5% w/v FeH_14_O_11_S·7H_2_O, 1% w/v (NH_4_)2MoO_4_, 1N H_2_SO_4_ aq.) was incubated for an hour. Samples were placed in quartz cuvettes and absorbance at 660 nm was measured using a plate spectrophotometer. All P*i* concentrations were calculated based on a standard curve from a set of standards made with known P*i* concentrations.

### RNA isolation and Quantitative Real-Time PCR

RNA was extracted from 10-day-old whole seedlings grown on 1/2X MS media containing 1% sucrose and 0.8% agar (Plant RNeasy kit (Qiagen)). During this process, samples were DNAse treated with RNase-free DNase (Qiagen). cDNA was generated using the iScript cDNA Synthesis Kit (Bio-Rad Laboratories, USA) using the manufacturer’s instructions. Quantitative PCR was performed using materials and methods previously described (Donahue *et al*., 2010). All primer sequences used in the qRT-PCR analysis can be found in Table S1. Relative expression was calculated using the ΔΔCT method. Expression data was normalized to *PEX4* and relative to the fold change in WT.

## ACCESSION NUMBERS

*ipk1* (SALK_065337; AT5G42810) WT Pennycress (Spring32-10 line; CS29085)

## ACKNOWLEDGMENTS

This work was funded in part by the USDA National Institute of Food and Agriculture, Hatch project VA-136334, the National Science Foundation to GG (MCB 1616038), and through the Virginia Tech Institute for Critical Technologies and Applied Sciences (ICALS). We gratefully thank Dr. John Sedbrook and Dr. Maliheh Esfahanian for their advice and expertise on how to grow and transform pennycress.

## SHORT LEGENDS FOR SUPPORTING INFORMATION

Supporting figures:

Figure S1. Additional InsP and PP-InsP profiles of DDP1 OX transgenics

Figure S2. DDP1-GFP localization in *Arabidopsis* epidermal cells

Figure S3. DDP1-GFP expression after 24 and 48 hours

Figure S4. *N. benthamiana* leaves co-infiltrated with YFP-DDP1, mCherry, and ER-mCherry

Figure S5. Transient DDP1-GFP expression in *N. benthamiana* leaves 48 hours post-infiltration

Figure S6. Homozygous pennycress DDP1-B OX plants

## Supporting tables

Table S1. Oligonucleotides used in this study

---

**Supplemental Table 1.**
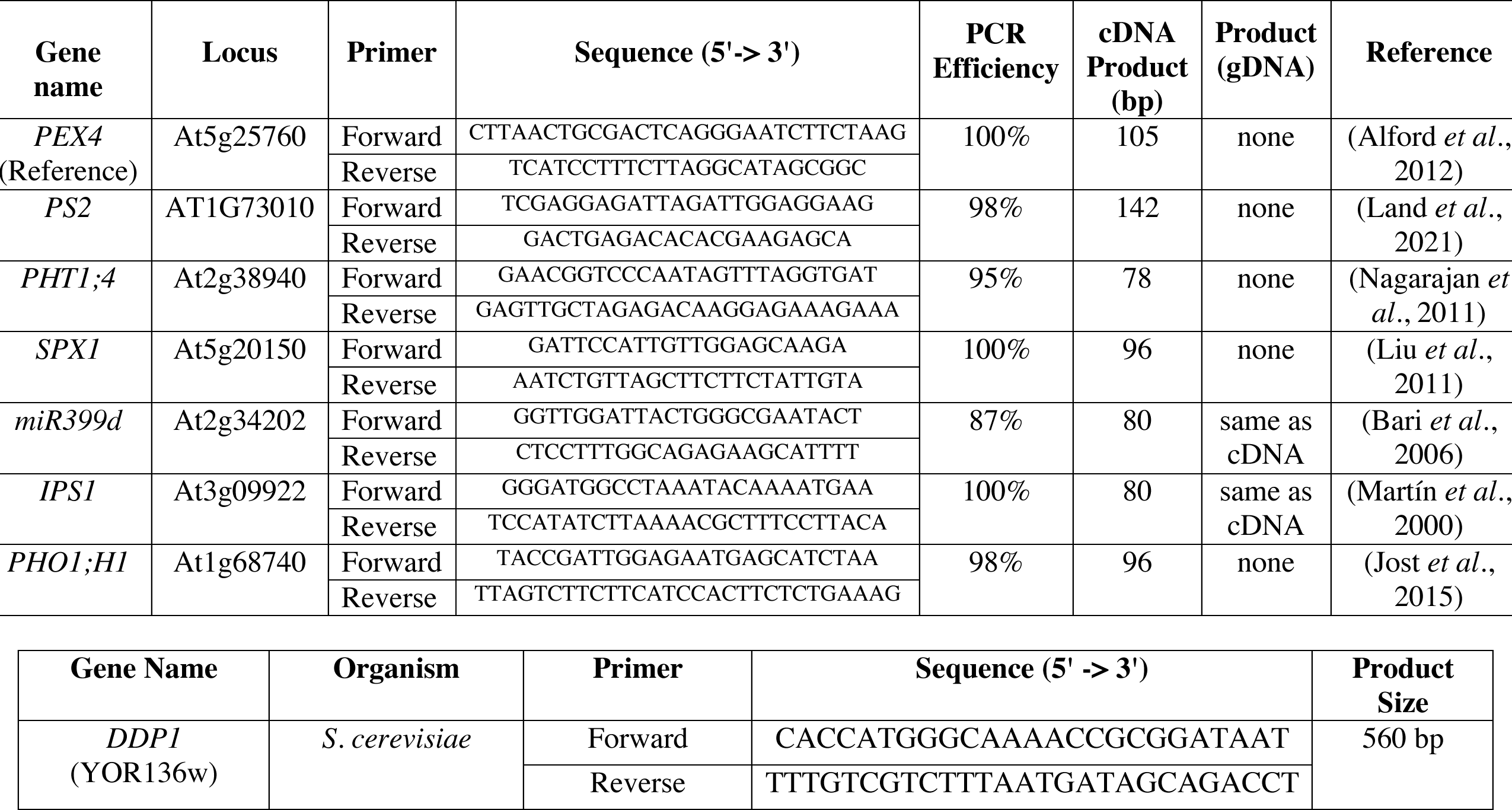
Oligonucleotide primers used in this study. PCR efficiencies for each primer pair shows the average of n = 3 technical replicates.

