## Supplemental Figures 1-6 for "Translational approach to increase phosphate accumulation in two plant species through perturbance of inositol pyrophosphates": SupplementalTextFile.txt

Supporting Figure S1. Additional InsP and PP-InsP profiles of DDP1 OX transgenics. (a) WT and DDP1 OX transgenics were grown for 14 days on semi-solid 1/2X MS media with 0.2% agar then 100 μCi [3H]-<i>myo</i>- inositol was added for 4 days. All InsPs were extracted and separated using anion exchange HPLC. (b) InsP<sub>8</sub>/InsP<sub>7</sub> ratios in DDP1 OX transgenics. Error bars, SD of n = 2-3 independent experiments. 


Supporting Figure S2. DDP1-GFP localization in <i>Arabidopsis</i> epidermal cells. Mature leaves of 3-week-old soil grown <i>Arabidopsis</i> were imaged. WT (a-c), DDP1-H (d-f), DDP1-I (g-i), and DDP1-A (j-l). Images (a, d, g, j) were imaged using the GFP channel and chlorophyll autofluorescence was imaged in images (b, e, h, k). All images are presented as maximum intensity projections from confocal Z-stack optical sections. All scale bars = 50 µm.

Supporting Figure S3. DDP1-GFP expression after 24 and 48 hours. DDP1-GFP was transiently expressed in <i>N. benthamiana</i> leaves and imaged 24 (a-c) and 48 (d-f) hours post-infiltration using confocal microscopy. Scale bar = 50 µm. Data in d-f is the same as presented Figure 3e. Images a and d were imaged using the GFP channel and chlorophyll autofluorescence was imaged in b and e. All images are presented as maximum intensity projections from confocal Z-stack optical sections. All scale bars = 50 µm.

Supporting Figure S4. <i>N. benthamiana</i> leaves co-infiltrated with YFP-DDP1 (a,d,g,j), mCherry (b,e,h), and ER-mCherry (k). Cells were imaged at 24 (a-c), 48 (d-f, j-l), and 72 (g-i) hours post-infiltration using confocal microscopy. (c,f,j) show the YFP-DDP1 + free mCherry merge and (l) shows the YFP-DDP1 + ER-mCherry merge. All images are presented as maximum intensity projections from confocal Z-stack optical sections. All scale bars = 50 µm.

Supporting Figure S5. Transient DDP1-GFP expression in <i>N. benthamiana</i> leaves 48 hours post-infiltration. DDP1-GFP (a,d,g,j) was co-expressed with free mCherry (b,e) and ER-mCherry (h,k). (c,f) show the DDP1-GFP + free mCherry merge and (i,l) show the DDP1-GFP + ER-mCherry merge. All images are presented as maximum intensity projections from confocal Z-stack optical sections. All scale bars = 50 µm.

Supporting Figure S6. Homozygous pennycress DDP1-B OX plants. Homozygous DDP1 overexpression severely stunted plant growth (a) and caused leaves to manifest detrimental amounts of leaf tip necrosis and chlorosis (b). 
