## Supplementary figures and images for "Translational approach to increase phosphate accumulation in two plant species through perturbance of inositol pyrophosphates"

### SFigure1.tiff

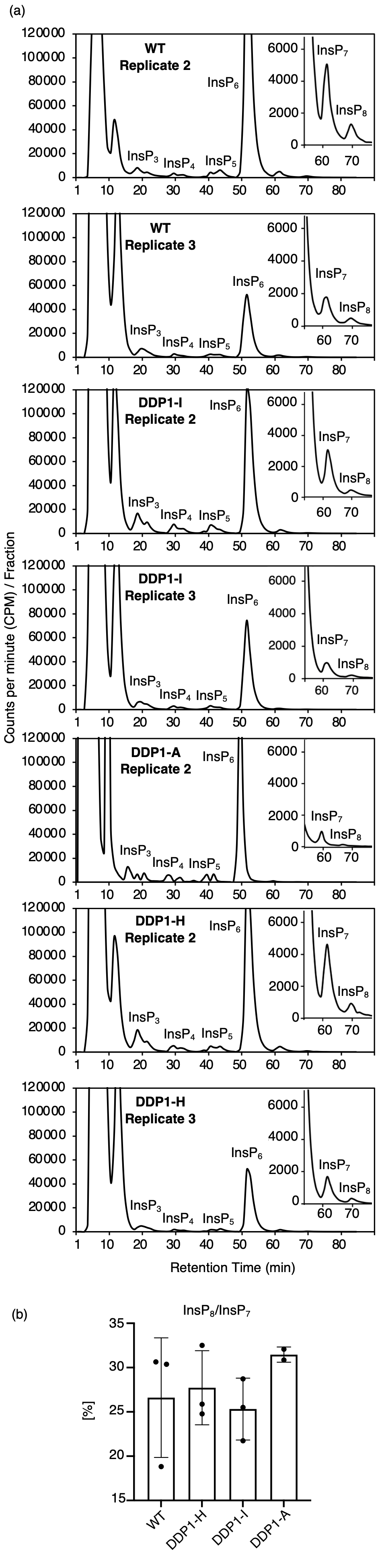

### SFigure2.tiff

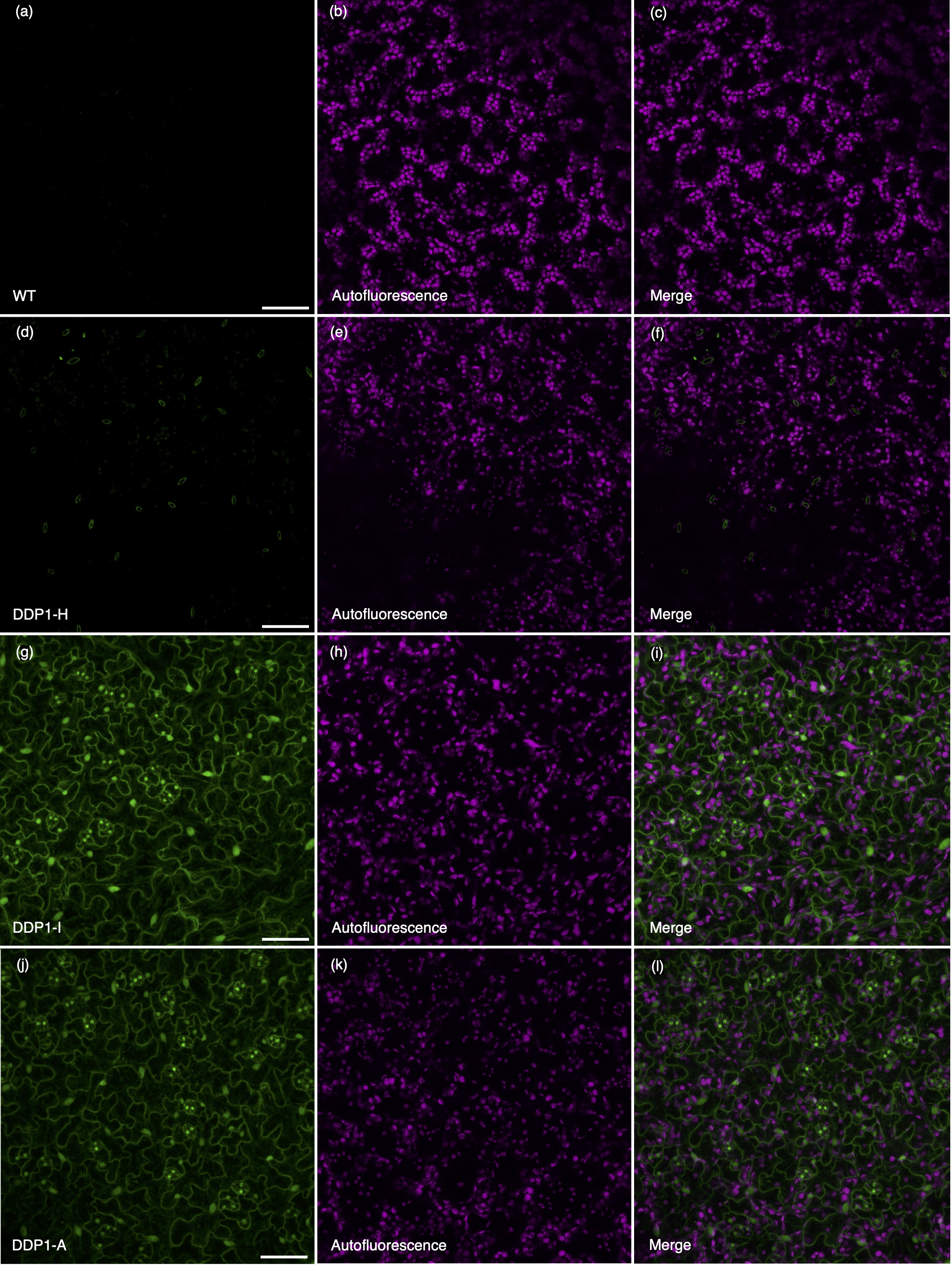

### SFigure3.tiff

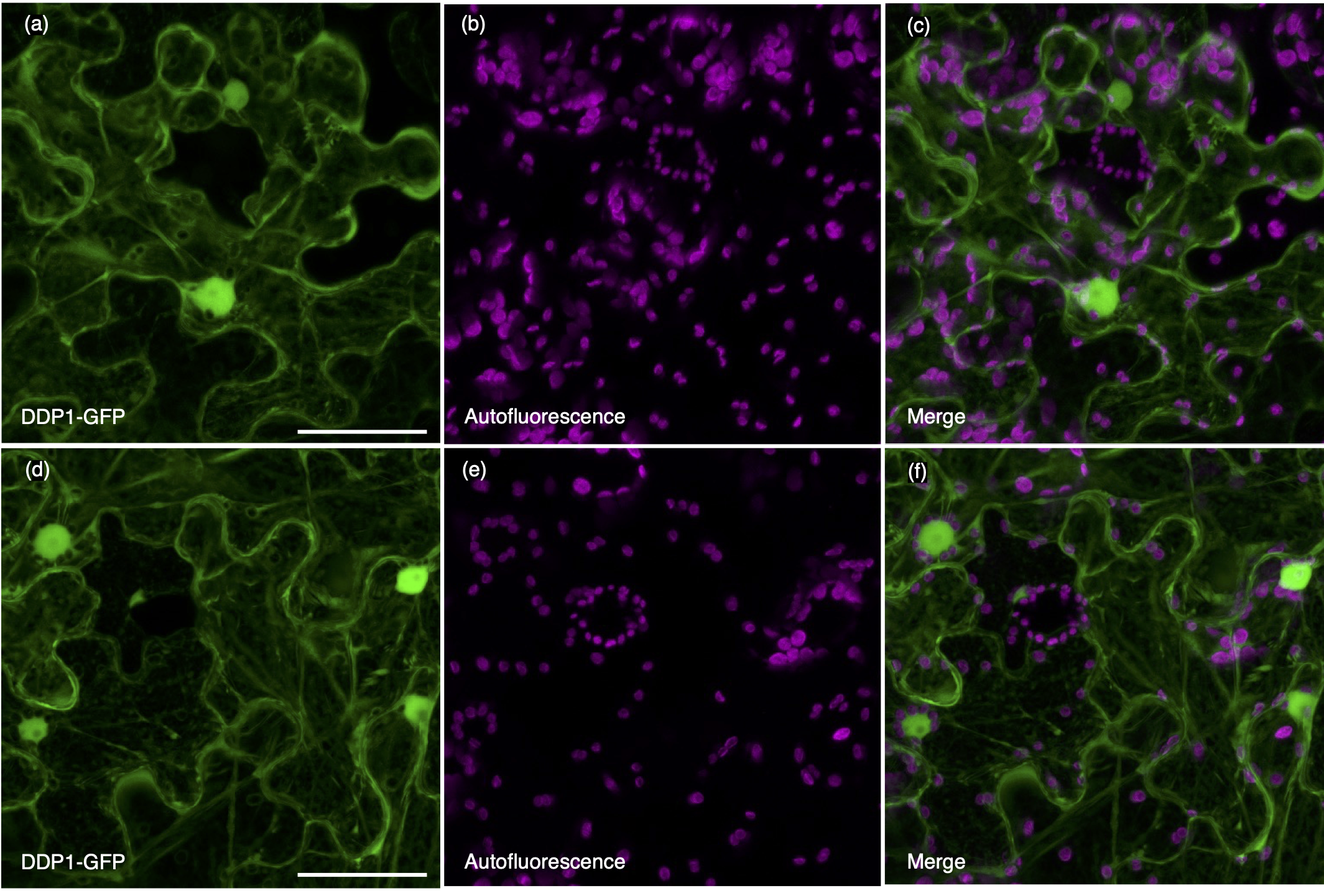

### SFigure4.tiff

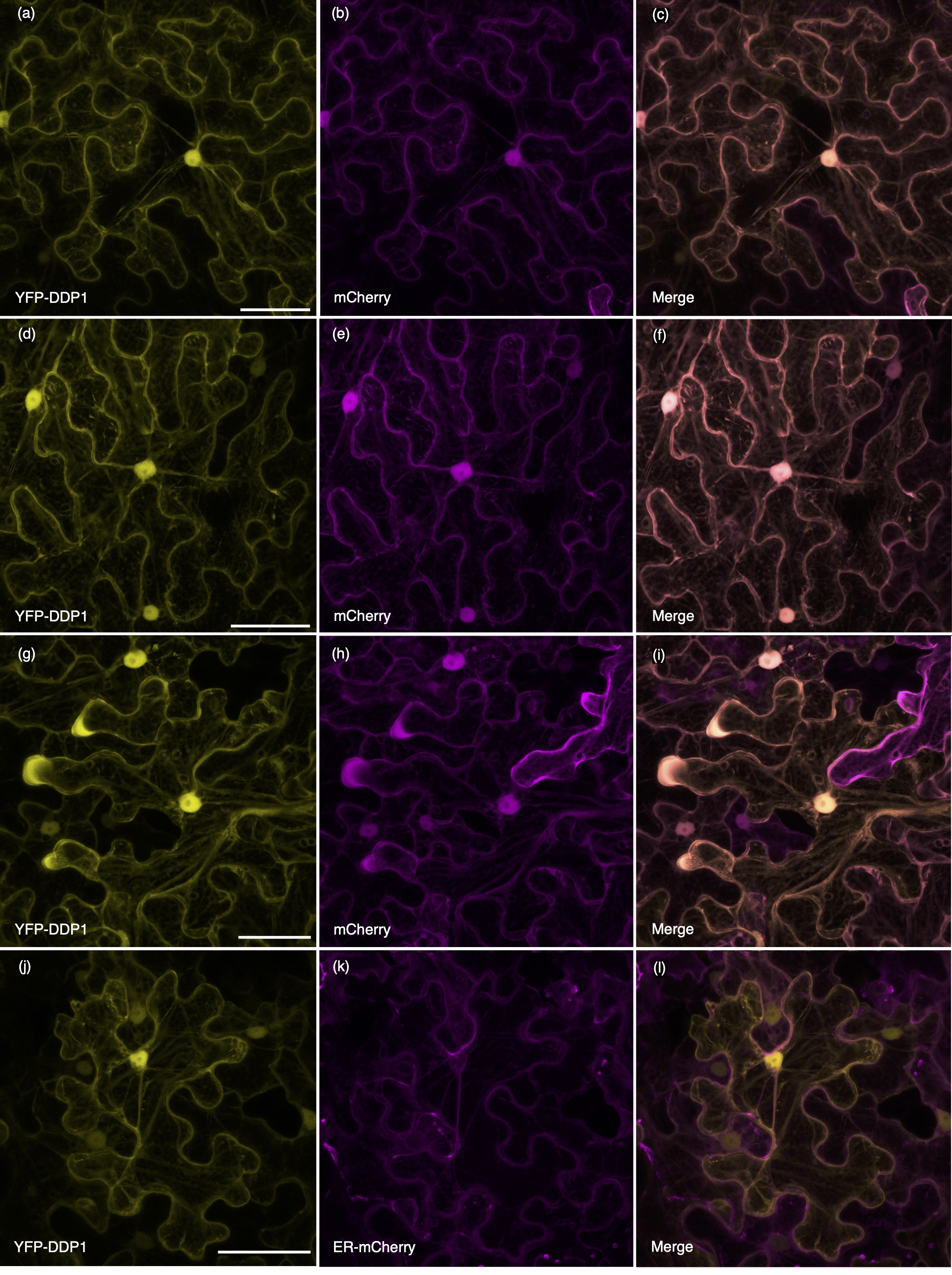

### SFigure5.tiff

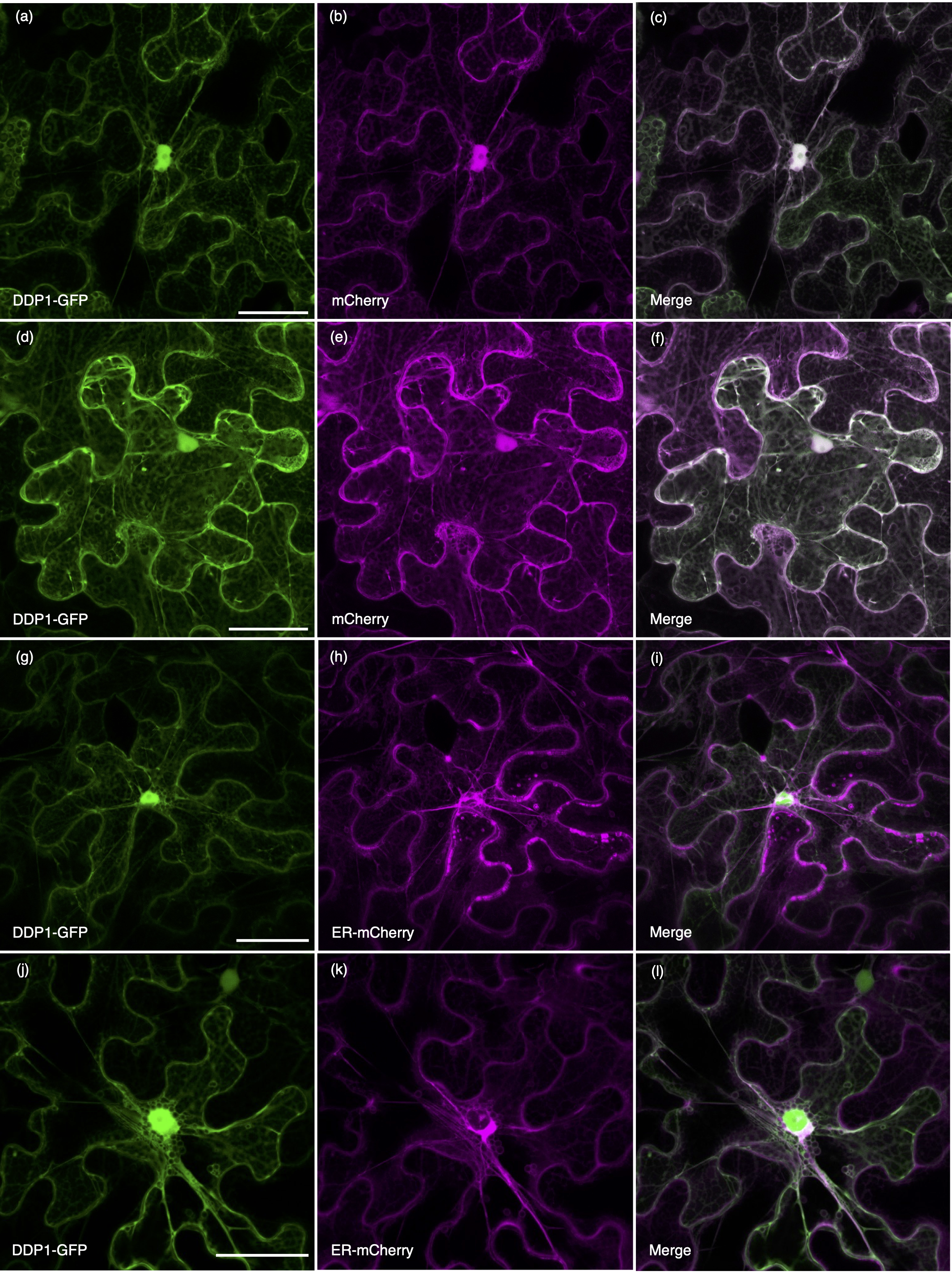

### SFigure6.tiff

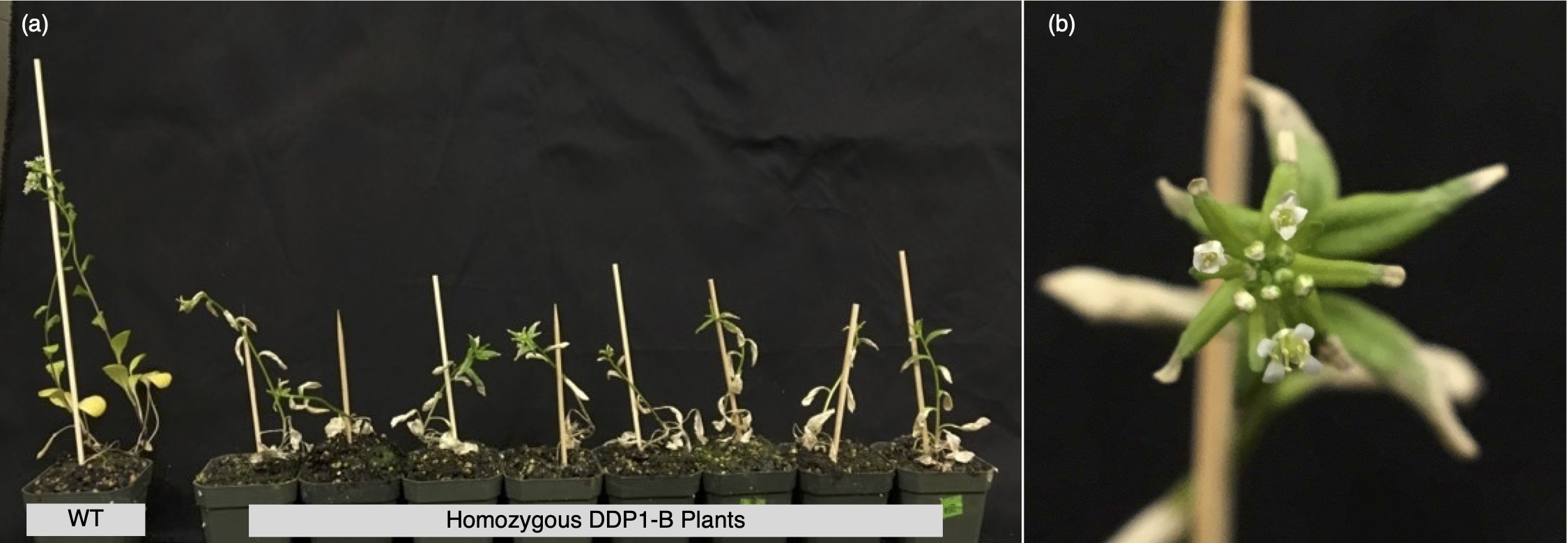
